# Deep multitask learning of gene risk for comorbid neurodevelopmental disorders

**DOI:** 10.1101/2020.06.13.150201

**Authors:** Ilayda Beyreli, Oguzhan Karakahya, A. Ercument Cicek

## Abstract

Autism Spectrum Disorder (ASD) and Intellectual Disability (ID) are comorbid neurodevelopmental disorders with complex genetic architectures. Despite large-scale sequencing studies only a fraction of the risk genes were identified for both. Here, we present a novel network-based gene risk prioritization algorithm named DeepND that performs cross-disorder analysis to improve prediction power by exploiting the comorbidity of ASD and ID via multitask learning. Our model leverages information from gene coexpression networks that model human brain development using graph convolutional neural networks and learns which spatio-temporal neurovelopmental windows are important for disorder etiologies. We show that our approach substantially improves the state-of-the-art prediction power in both single-disorder and cross-disorder settings. DeepND identifies prefrontal and primary motor-somatosensory cortex brain region, and periods from early fetal to mid fetal periods and from early childhood to young adulthood as the highest neurodevelopmental risk windows for both ASD and ID. Finally, we investigate frequent ASD and ID associated copy number variation regions and report our findings for several susceptibility gene candidates. DeepND can be generalized to analyze any combinations of comorbid disorders and is released at http://github.com/ciceklab/deepnd.

## 1 Introduction

Autism Spectrum Disorder (ASD) is a common neurodevelopmental disorder with a complex genetic architecture in which around a thousand risk genes have a role [36]. Large consortia efforts have been paving the way for understanding the genetic, functional and cellular aspects of this complex disorder via large scale exome [18, 41, 73, 67, 66, 62, 40] and genome [33, 23, 20, 57, 1, 3] sequencing studies. Latest and also the most comprehensive study to date analyzed *∼* 36*k* samples (6, 430 trios) to pinpoint 102 risk genes (*FDR* ≤ 0.1) [75]. Overwhelming evidence suggests that genetic architectures of neuropsychiatric disorders overlap [60, 71, 53]. For instance, out of the twenty five SFARI Cat I ASD risk genes (i.e., highest risk), only five are solely associated with ASD. Genes like *CHD2, SCN2A* and *ARID1B* are associated with six neurodevelopmental disorders. Intellectual Disability (ID) is one of such comorbid disorders which manifests itself with impaired mental capabilities. Reminiscent of ASD, ID also has a complex genetic background with hundreds of risk genes involved and identified by rare *de novo* disruptive mutations observed in whole exome and genome sequencing studies [17, 24, 26, 69, 77, 80, 86]. ASD and ID are frequently observed together [61]. In 2018, CDC reported that 31% of children with ASD were also diagnosed with ID and 25% were borderline [5]. They also share a large number of risk genes [54]. Despite these similarities, Robinson et al. also point to differences in genetic architectures and report that intelligence quotient (IQ) positively correlates with family history of psychiatric disease and negatively correlates with *de novo* disruptive mutations [70]. Yet, the shared functional circuitry behind is mostly unknown.

The current lack of understanding on how comorbid neuropsychiatric disorders relate mostly stems from the incomplete profiling of individual genetic architectures. Statistical methods have been used to assess gene risk using excess genetic burden from case-control and family studies [36] which are recently extended to work with multiple traits [64]. Yet, these tools work with genes with observed disruptive mutations (mainly *de novo*). It is often of interest to use these as prior risk and obtain a posterior gene interaction network-adjusted risk which can also assess risk for genes with no prior signal. Network-based computational gene risk prediction methods come handy for (i) imputing the insufficient statistical signal and providing a genome-wide risk ranking, and (ii) finding out the affected cellular circuitries such as pathways and networks of genes [36, 30, 25, 38, 65, 28, 27, 63, 8, 21]. While these methods have helped unraveling the underlying mechanisms, they have several limitations. First, by design, they are limited to work with a single disorder. In order to compare and contrast comorbid disorders such as ASD and ID using these tools, one approach is to bag the mutational burden observed for each disorder assuming two are the same. However, disorder specific features are lost as a consequence [30]. The more common approach is to perform independent analyses per disorder and intersect the results. Unfortunately, this approach ignores valuable source of information coming from the shared genetic architecture and lose prediction power as per-disorder analyses have less input (i.e., samples, mutation counts) and less statistical power [75, 10, 38]. Second, current network-based gene discovery methods can work with one or two integrated gene interaction networks [28, 56, 38, 49, 8]. This means numerous functional interaction networks (e.g., co-expression, protein interaction etc.) are disregarded which limits and biases the predictions. Gene co-expression networks that model brain development are a promising source of diverse information regarding gene risk, but currently cannot be fully utilized, as the signal coming from different networks cannot be deconvoluted. Usually, investigating such risky neurodevelopmental windows is an independent downstream analysis [83, 49]. Should this process be integrated within the risk assessment framework, it has potential to provide valuable biological insights and also to improve the performance of the genome-wide risk assessment task.

Here, we address these challenges with a novel cross-disorder gene discovery algorithm (Deep Neurodevelopmental Disorders algorithm - *DeepND*.) For the first time, DeepND analyzes comorbid neurodevelopmental disorders simultaneously over multiple gene co-expression networks and explicitly learns the shared and disorder-specific features using multitask learning. Thus, the predictions for the disorders *depend* on each other’s genetic architecture. The proposed DeepND architecture uses graph convolution to extract associations between genes from gene co-expression networks. This information is processed by a mixture-of-experts model that can self-learn critical neurodevelopmental time windows and brain regions for each disorder etiology which makes the model interpretable. We provide a genome-wide risk ranking for each disorder and show that the prediction power is improved in both singletask (single disorder) and multitask settings. DeepND identifies prefrontal cortex brain region and from early fetal to late mid-fetal and from mid-childhood to young adulthood periods as the highest neurodevelopmental risk windows for both disorders.

Finally, we investigate frequent ASD and ID associated copy number variation regions and pinpoint risk genes that are disregarded by other algorithms. The software is released at http://github.com/ciceklab/deepnd. This neural network architecture can easily be generalized to other disorders with a shared genetic component and can be used to prioritize focused functional studies and possible drug targets.

## 2 Results

Using a deep learning framework which combines graph convolutional neural networks with a mixture of experts model, we perform a genome-wide risk assessment for ASD and ID simultaneously in a multitask learning setting and detect neurodevelopmental windows that are informative for risk assessment for each disorder. Our results point to the shared disrupted functionalities and novel risk genes which provides a road map to researchers who would like to understand the ties between these two comorbid disorders.

### 2.1 Genome-wide Risk Prediction for ASD and ID

To have a model of evolving gene-interactions throughout brain development, we extracted 52 gene co-expression networks from the BrainSpan dataset [78, 44] using hierarchical clustering of brain regions and a sliding window approach [83]. These networks are used to assess the network-adjusted posterior risk for each gene given disease agnostic the prior risk features such as pLI. Note that one can use disease specific features such as mutation burden in case-control or family studies. However, current knowledge on ground truth risk genes mostly rely on this data and using these as features would cause a leak in the training procedure.

The deep neural network architecture (DeepND) performs a semi-supervised learning task, which means a set of positively and negatively labeled genes are required as ground truth per disorder. As also done in [49], we obtained 594 ASD-positive genes (Supplementary Table 1) from SFARI dataset (http://gene.sfari.org) [2] which are categorized into 3 evidence levels based on the strength of evidence in the literature. For ID, we curated a ground truth ID risk gene set of 580 genes using landmark review studies on ID gene risk [26, 34, 79, 39, 12, 45] (Supplementary Table 2). We generated three evidence level sets similar to the ASD counterpart: E1, E2 and E3 sets where each set includes genes which are recurrently indicated in multiple studies. As for the negatively labeled genes, we use 1074 non-mental-health related genes for both disorders which is curated by Krishnan et al., (2016) after discarding 181 of them which are shown to be related to other neurodevelopmental disorders, such as epilepsy, schizophrenia or bipolar disorder (Supplementary Table 1).

DeepND uses the multitask learning paradigm where multiple tasks are solved concurrently (i.e., genome wide risk assessment for ASD and ID). Thus, the network learns a shared set of weights for both disorders and also disorder-specific set of weights (Figure 1). First, the model inputs pLI for each gene and using fully-connected layers produces an embedded feature set per gene. The set of weights learnt in these layers are affected by the ground truth labels for both disorders, and thus, are shared. Then, the architecture branches out to 2 single task layers, one per disorder (blue for ASD and yellow for ID in Figure 1). For each single task branch, these embeddings are input to 52 graph-convolutional neural networks (GCNs). Each GCN processes a co-expression network that represents a neurodevelopmental window and extracts network-adjusted gene risk signatures (i.e., embeddings) [75, 63] (Methods). Finally, these embeddings are fed into a fully-connected gating network along with the prior risk features. The gating network assigns a weight to each GCN which is proportional to the informativeness of the embedding coming from each neurodevelopmental window. Thus, the model also learns which windows are important for prediction of and ASD/ID risk genes. This also means they are important in the etiology of the disorder. In the end, each disorder-specific subnetwork produces a genome-wide ranking of genes being associated with that disorder along with risk probabilities. To quantify the contribution of the co-analyzing comorbid disorders (i.e., multitask) as opposed to individual analysis (i.e., singletask), we also present our results of DeepND when it is run on a single task mode (DeepND-ST). In this mode, the fully connected layers that generates embeddings are removed and the feature (i.e., pLI) is directly fed into the GCNs (Methods). We show that the genome-wide risk assessment of DeepND is robust, precise and sensitive and substantially improves the state of the art both in terms of performance and interpretability of the predictions.

**Fig. 1:**
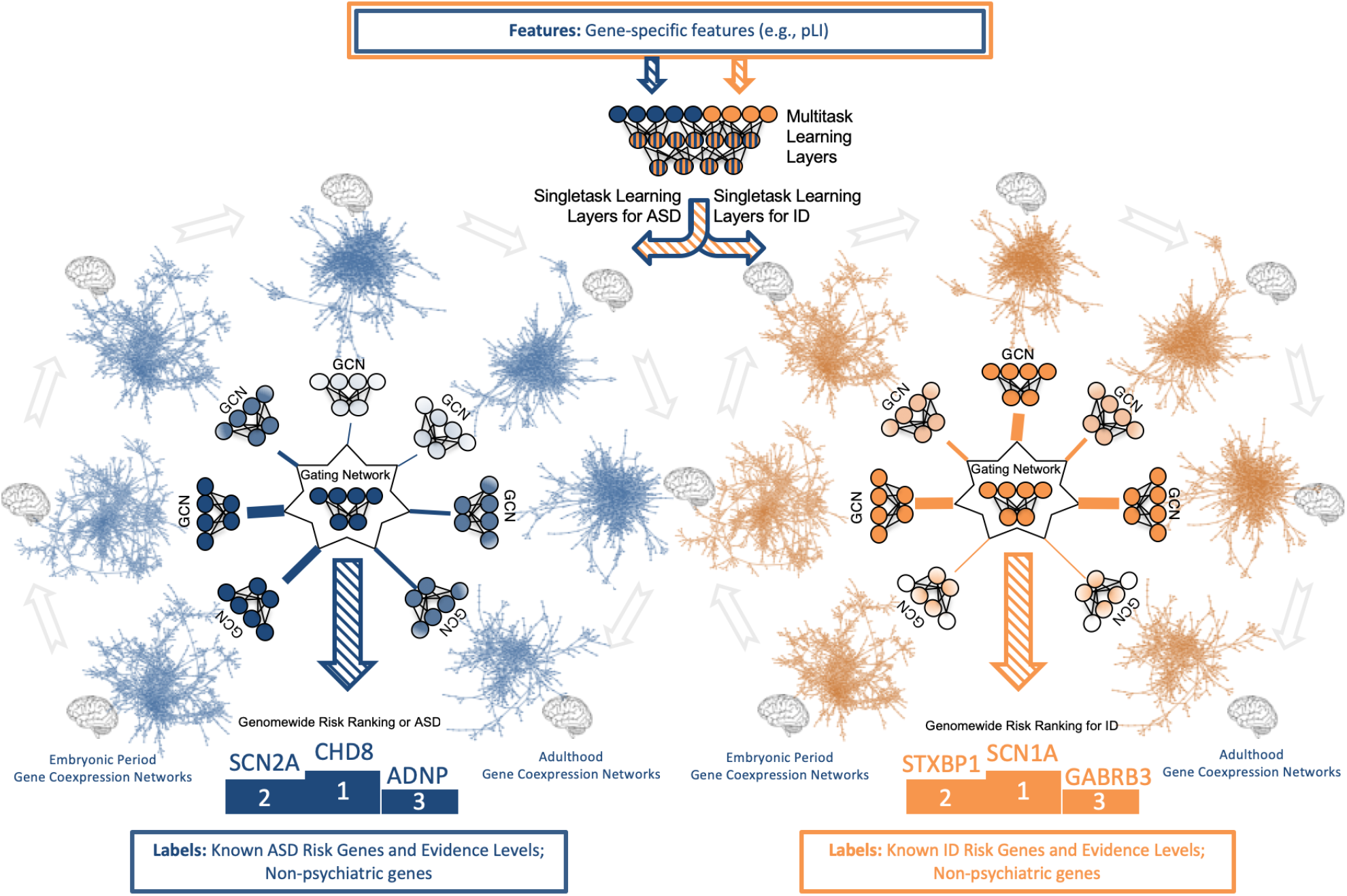
System model of the proposed deep learning architecture for genome-wide cross-disorder risk assessment (DeepND). The algorithm takes the following information as input: (i) gene specific features such as pLI; (ii) disorder specific ground truth genes that are labeled as positive with varying level of evidence based on a literature search; and (iii) non-psychiatric genes which are labeled as negative. The features are passed through fully-connected multitask layers that learn shared weights for both disorders and produces a new feature representation. This new representation is then input to singletask graph convolutional neural networks (GCNs), each processing one of fifty-two gene co-expression networks that represent different brain regions and neurodevelopmental time windows. The output of GCNs are then weighted by the Gating Network to learn which networks are informative for the gene risk assessment (shade of the network indicates importance). Thus, DeepND learns which neurodevelopmental windows confer more risk for each disorder’s etiology. The final output is a genome-wide risk probability ranking per disorder, which are then used for various downstream analyses to understand the underlying functional mechanisms and to compare/contrast both disorders. The singletask layers are exclusively trained with the ground truth genes of the disorder they belong. Thus, they learn only disorder specific parameters and disorderspecific networks that implicate risk.

We benchmark the performances of DeepND and other gene discovery algorithms for neurodevelopmental disorders. First, we compare DeepND and the state of the art algorithm by Krishnan et al. using their experimental settings. Evaluating performances of the algorithms using E1 genes through 5-fold cross shows that DeepND achieves a median AUC of 94% and a median AUPR of 83% for ASD, which correspond to 4% and 47% improvements over the evidence weighted SVM algorithm of Krishnan et al., respectively (Figures 2a and 2b). Similarly for ID, DeepND achieves a median AUC of 87% and a median AUPR of 57%, which corresponds to an improvement of 15% and 41%, respectively. We observe that even the singletask mode DeepND-ST performs better than Krishnan et al.’s algorithm in all settings other than ID AUPR. As expected, the multitask setting of DeepND performs better than DeepND-ST and leads to improved AUC (up to 13%) and AUPR (up to 20%) for both disorders.

**Fig. 2:**
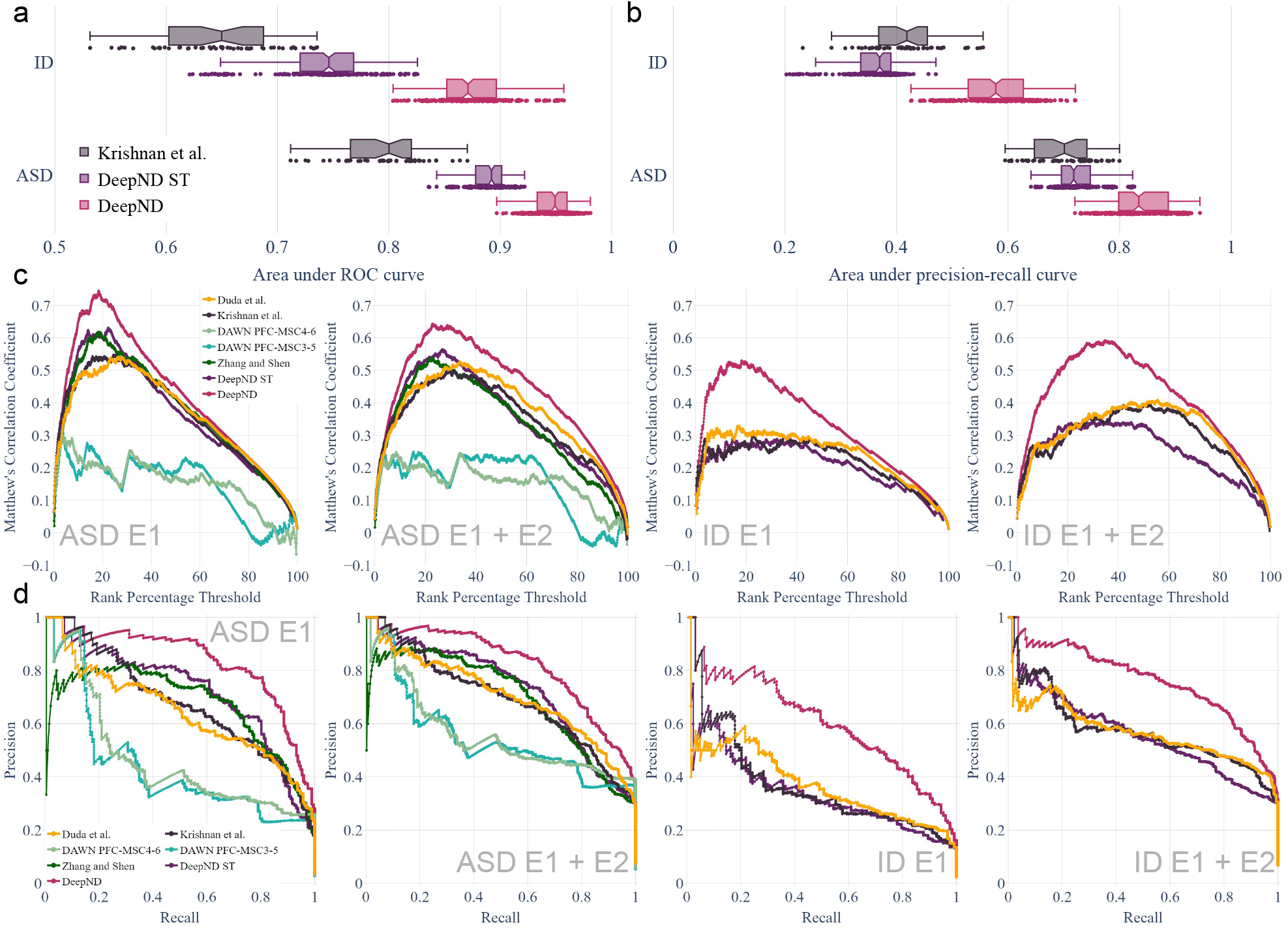
Evaluation of DeepND genome-wide risk assessment for ASD and ID. (**a**) The area under ROC curve distributions of Krishnan et al., DeepND and DeepND-ST for ASD and ID genome-wide risk assessments. Every point corresponds to the performance on a test fold in the repeated cross validation setting (**b**) The area under precision-recall curve distributions for the same methods as in (**a**). Center line: median; box limits: upper and lower quartiles; whiskers: 1.5x interquartile range; points: outliers. (**c**) MCC between the rankings of each method and the ground truth genes are shown for varying rank percentage threshold values (x axis). This ranking threshold p sets the top p% of a ranking as positive predictions. Methods compared are DeepND, DeepND - ST, Evidence weighted SVM of Krishnan et al., DAWN (only compared for ASD, due to low intersection ratio with ground truth gene set of ID), DAMAGES score of Zhang and Shen, and Random Forest based method of Duda et al.. Results for ASD and ID shown when (i) E1 genes are used as the true risk genes and (ii) E1+E2 genes are considered as the true risk genes. (**d**) Precision - Recall curves to compare the same set of methods in panel c).

We also compare the ranking performance DeepND with the following methods from the literature using a rank percentage threshold based method: Krishnan et al.[49], DAWN [56], DAMAGES score [84], and Random Forest based method of Duda et al. [21]. That is, we obtain the final ranking of each method and mark *p*% of the top genes predicted as risk genes. We calculate the Matthews Correlation Coefficient of these marked lists with the ground truth. We vary *p* by 1% and obtain the plots in Figure 2c. Results show that for varying threshold levels, DeepND curve always dominates all others indicating that it achieves the highest correlation with the ground truth both when we use E1 and E1 + E2 genes as positive ground truth. Comparing precision-recall performance of the same set of methods show that the PR curve of DeepND domiates all other in all settings and for both disorders (Figure 2d).

Finally, we compare DeepND with an ensemble learner, forecASD [8]. This method uses the outputs of other methods including the ones we compared above and learns to predict ASD/ID risk genes in a semi-supervised setting. However, labels used by forecASD are also used by the voter methods whose outputs are used as features, which suggests information leakage during the training process and the performance might be overly optimistic. Comparing DeepND and forecASD shows that despite this advantage of forecASD, DeepND performs slightly worse or on par with respect to rank-percentage threshold curve and PR curve comparisons in ASD (see Supplementary Figure 1). DeepND performs better than forecASD in all settings for ID. We also considered adding DeepND as an additional voter into the ensemble of forecASD to check if it benefits the final performance. For both disorders and in all settings forecASD that uses DeepND as a voter outperforms both DeepND and original forecASD. This shows the informativeness of DeepND predictions.

### 2.2 Critical neurodevelopmental windows for ASD and ID Risk

We observe that the top percentile genes in the DeepND ASD and ID rankings have a high overlap with a Jaccard index of 65%. Top 3 deciles also have relatively high overlap with Jaccard indices 0.66, 0.71, and 0.69, respectively (Supplementary Figure 2).

To further investigate the shared genetic component, we focus on the spatio-temporal neurodevelopmental windows that are deemed important by DeepND for accurate ranking of risk genes for both disorders. The neural network analyzes co-expression networks that represent 13 neurodevelopmental time windows (from embryonic period to late adulthood; Figure 3a) and 4 brain region clusters (PFC-MSC: Prefrontal and motor-somatosensory cortex; MDCBC: Mediodorsal nucleus of the thalamus and cerebellar cortex; V1C-STC: Primary visual and superior temporal cortex; SHA: Striatum, Hippocampus, Amygdala) generated using Brainspan RNA-Seq dataset [44] in accordance with Willsey et al. [83] (Methods; Figure 3b).

**Fig. 3:**
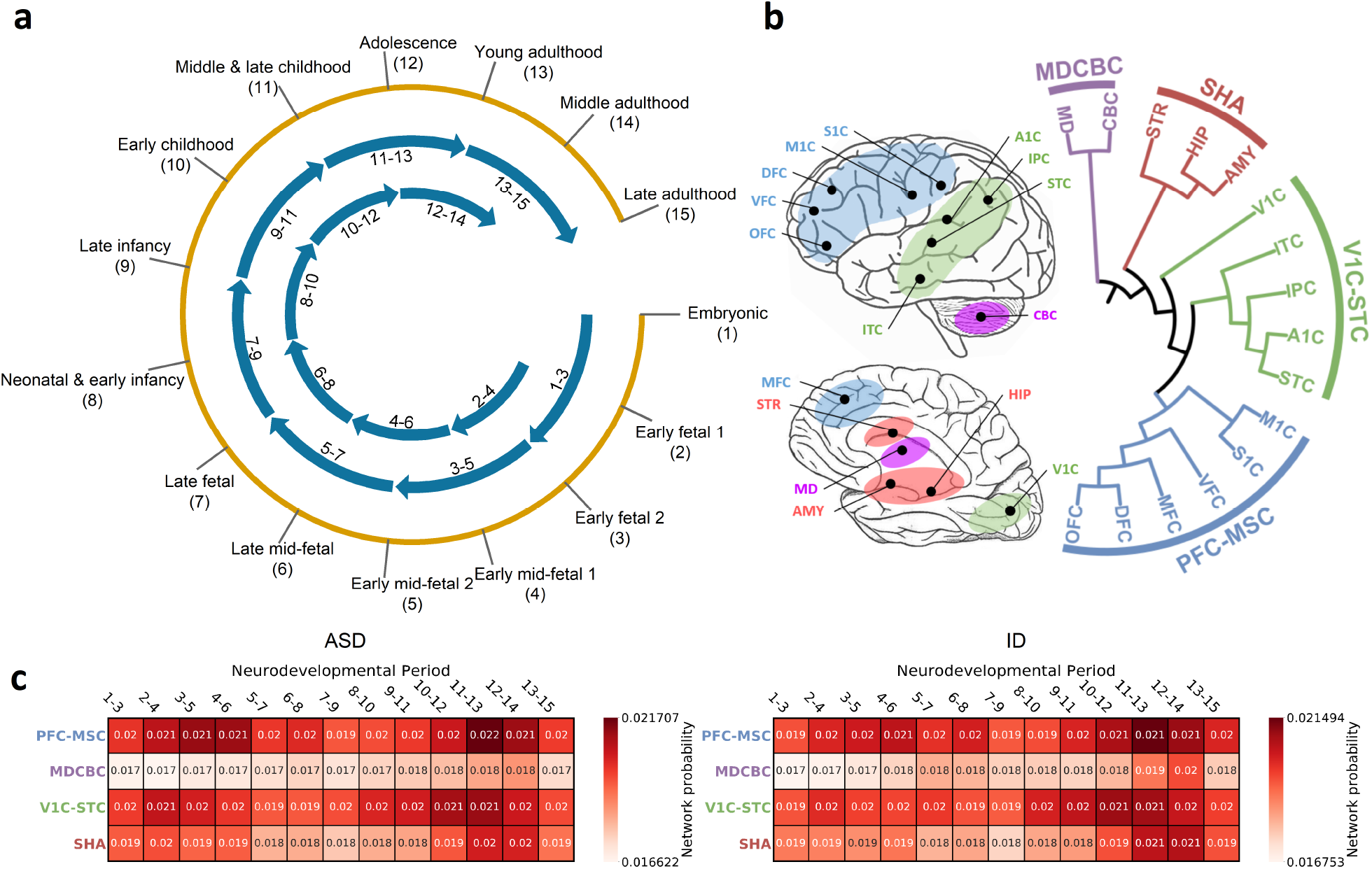
The BrainSpan dataset which models the spatio-temporal gene expression of human neurodevelopment [44] is used to obtain gene co-expression networks. This dataset contains samples from 12 brain regions and spans 15 time points from early fetal period to late adulthood. **(a)** We generate 13 neurodevelopmental time-windows using a sliding window of length 3 which provides sufficient data in each window as also done by Willsey et al. [83]. **(b)** We obtain 4 brain region clusters based on transcriptional similarity during fetal development which also reflects topographical closeness and functional segregation. The brain regions considered are as follows. HIP: Hippocampal anlage (for (1)–(2)), Hippocampus (for (3)–(15)); OFC: Orbital prefrontal cortex; DFC: Dorsal prefrontal cortex; VFC: Ventral prefrontal cortex; MFC: Medial prefrontal cortex; M1C: Primary motor cortex; S1C: Primary somatosensory cortex; IPC: Posterior inferior parietal cortex; A1C: Primary auditory cortex; STC: Superior temporal cortex; ITC: Inferior temporal cortex; V1C: Primary visual cortex; AMY: Amygdala; STR: Striatum; MD: Mediodorsal nucleus of the thalamus; CBC: cerebellar cortex. **(c)** Heatmaps show which spatio-temporal windows lead to assignment of higher risk probabilities to top percentile genes for ASD (left) and ID (right). The numbers in boxes are softmaxed outputs of each respective GCN, averaged for top percentile genes and then normalized. The weights assigned to each co-expression network by the MoE lets each GCN learn to make a better prediction.

We investigate using which neurodevelopmental windows more confidently distinguish disorder risk genes. Figure 3c shows normalized average probabilities assigned to top percentile risk genes (Methods). We observe that the networks of the PFC-MSC brain region, spanning several time windows from early fetal to mid fetal periods and from early childhood to young adulthood periods consistently are better predictors for ASD and ID risk. The periods 3-5 and 4-6 were also previously indicated as one of the highest risk regions and were subject to network analyses for ASD gene discovery [83, 56]. DeepND also captures strong signals from these windows, in addition to period 11-13, for both ASD and ID. We see that V1C-STC brain regions are second to PFC-MSC and MD-CBC regions provide the most subtle signal. We assess the empirical of significance of these findings via a permutation test. That is, we run DeepND in the exact same settings but shuffle the labels of the risk genes, keeping the same number of positively and negatively labeled genes. For each of the squares in Figure 3c, we obtain a distribution of average probabilities (background distribution) and check if the actual informativeness of that window is higher than the average background value. See these distributions in Supplementary Figures 4 and 5, for ASD and ID, respectively. We observe that informativeness of PFC-MSC windows are the most significant compared other brain regions and MD-CBC results are consistently insignificant. Overall, we observe that not only earlier time windows but also late time windows are important for both etiologies. Supplementary Figure 3 shows the heatmaps of DeepND-ST for ASD and ID.

The weakest source of information is MDCBC 2-4 network. We observe roughly 12.5k links between top percentile ASD and ID genes. The network contains close to 37.5m links. On the other hand, PFC-MSC 4-6 contains close to 785k edges. Yet, there exists close to 13k links between top percentile ASD and ID genes which is the reason behind DeepND focusing on this window as its top predictor. Visualization of the top 30 genes for each disorder in PFC-MSC 4-6 network with only very high co-expression links (*r*^2^ *>* 0.80) is provided in Figure 4a.

**Fig. 4:**
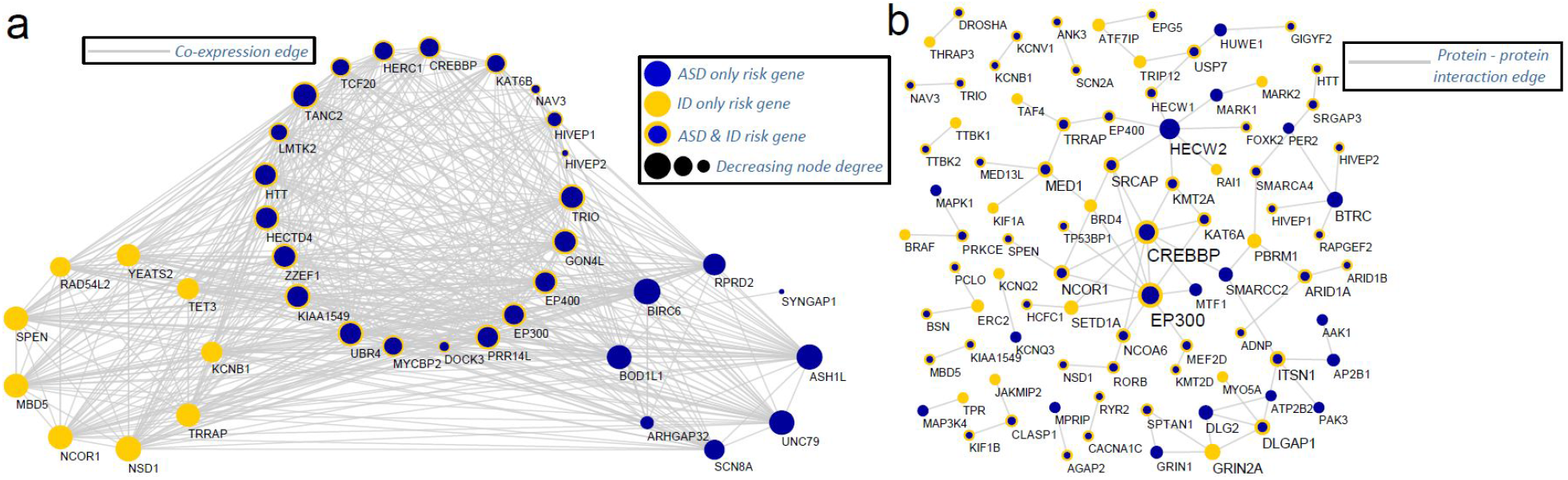
Network Analyses of the risk genes. (a) The co-expression relationships between top 30 ASD and ID risk genes in PFC-MSC 4-6 region. This is found as the informative co-expression network for DeepND and also indicated in the literature as an important window for ASD [83]. Only links for at least 0.95 absolute correlation and genes with at least one connection are shown. (b) The protein-protein interactions between the top percentile risk genes are shown. The PPIs are obtained from the tissue specific PPI network of frontal cortex in DifferentialNet database [6] Only interactions between the top percentile risk genes are shown.

### 2.3 Enrichment Analysis of the Predicted Risk Genes

In addition to the prediction performance benchmark above, we also evaluate the enrichment of our ASD and ID gene risk rankings in gene lists which are shown to be related to these disorders. That is, while these gene sets are not *ground truth* sets, they have been implicated as being associated with the etiology of either disorder. Thus, enrichment of members of these sets in the higher deciles of the genome-wide risk ranking of DeepND is an indication of the wellness of the ranking and also provides a means of comparing and contrasting the disrupted circuitries affected by ASD and ID. These lists are (i) targets of transcription regulators like CHD8 [15], FMRP [16, 4, 18], RBFOX [81, 82, 18] and TOP1 [46]; (ii) Susceptibility pathways like WNT signaling [43] and MAPK signaling [68]; and (iii) affected biological processes and molecular functions like, post-synaptic density complex [7, 87], histone modification [18, 42]).

The first decile of ASD-risk genes has the highest enrichment in all categories. All enrichments are significant with respect to Binomial test (Figure 5; Methods). We observe the same trend for ID that the top decile of the genome-wide risk ranking is the most enriched in most categories but the *P* values are more subtle. *CHD*8 (Chromodomain Helicase DNA Binding Protein 8) is the highest ASD-risk gene known to date with the highest mutation burden in large ASD cohorts [18, 75] and with downstream functional analysis [15]. Accordingly, DeepND ranks *CHD*8 as the 37^*th*^ top risk gene whereas Krishnan et al. ranks it 1943^*th*^ genome-wide. While it has a solid association to ASD etiology, the ties to ID is not well-established. Bernier et al. reports that 9 out of 15 ASD probands with mutations in *CHD*8 also has ID. In accordance, DeepND places CHD8 as the 53^*rd*^ highest risk gene for ID.

**Fig. 5:**
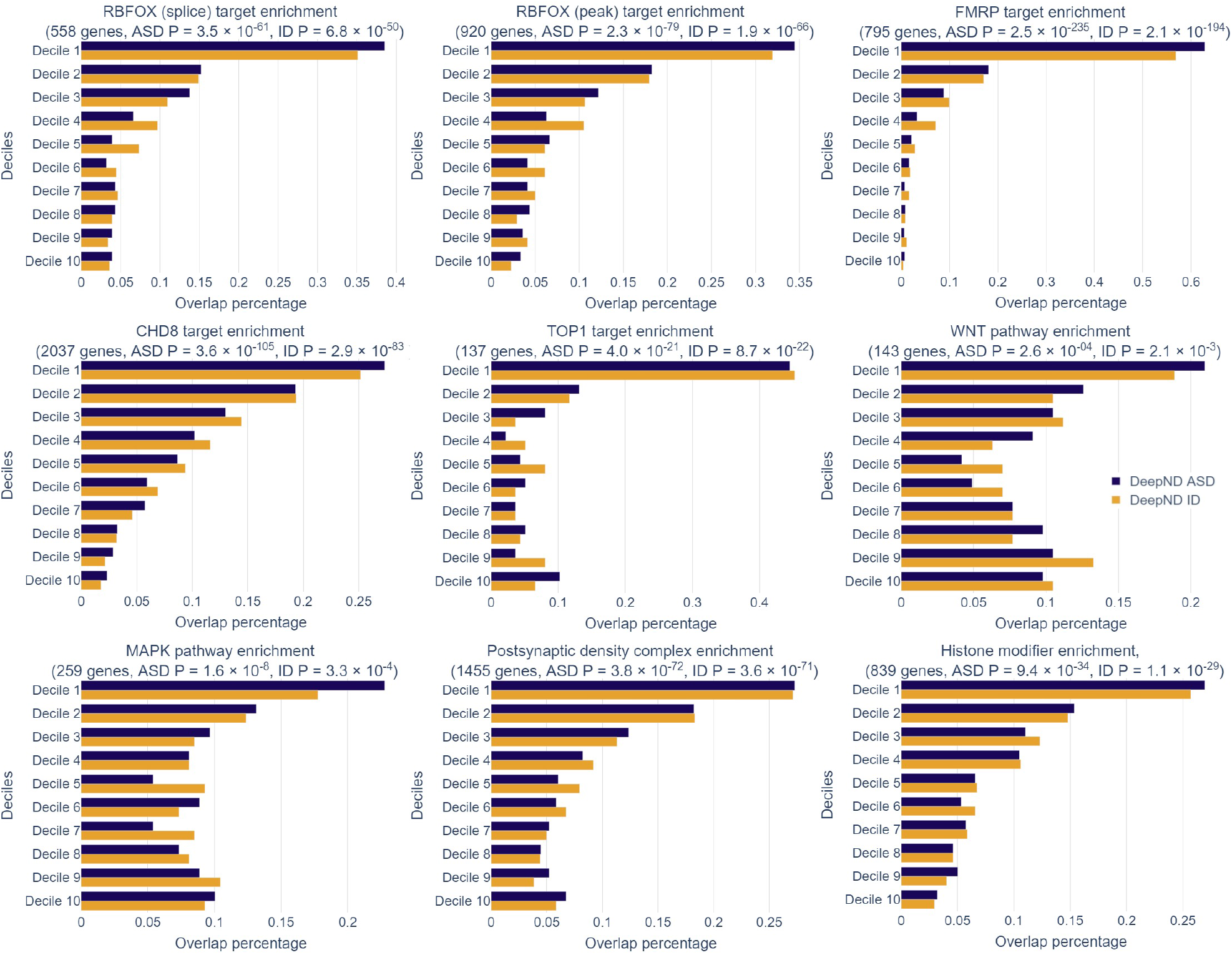
The enrichment of our ASD and ID gene risk rankings in various disease-related gene lists (i.e., each panel) are shown: (i) ASD and/or ID-related transcription regulators, (ii) ASD and/or ID-related pathways, and (iii) ASD and/or IDrelated biological functions or protein complexes. While these do not fully contain ground truth genes, they have been indicated in the literature as being enriched with risk genes for either disorder. Percentage of genes in the corresponding gene set (x axis) that occurred within each decile of the genome-wide risk ranking per ASD (blue) and ID (yellow) are shown. The gene sets used are as follows: (i) Targets of RBFOX (splice), (ii) Targets of RBFOX (splice target), (iii) Targets of FMRP (all peak), (iv) Targets of CHD8, (v) Targets of TOP1, (vi) WNT Pathway, (vii) MAPK Signaling Pathway, (viii) GTPase regulator activity, (x) Postsynaptic density complex genes, (xi) Synaptic genes (xii) Histone modifier genes.

We also perform an untargeted enrichment analysis of the top percentile predictions using the Enrichr system [11, 51]. Unsurprisingly, we find that for both disorders *nervous system development* is the top enriched Biological Process GO term (Fisher’s exact test, *P =* 1.48 × 10^*−*12^ for ASD and *P =* 3.95 × 10^*−*14^ for ID) which indicates that it is a shared function that is disrupted in the etiology of both disorders (Supplementary Table 3). As for the GO Molecular Function enrichment, two disorders overall have similar terms but top terms are different. The top function affected for ASD is *histone-lysine N-methyltransferase activity* indicating a disruption in the transcription factor activity whereas for ID the top affected function is *voltage-gated cation channel activity* which indicates a disruprion in synaptic activity.

As transcription factor activity is highly affected in both disorders, we further investigate if there are any master transcription regulators upstream that regulate the high risk genes for ASD and ID in the ChEA Database [52]. We find that SMARCD1 is the top transcription factor with 82 of its targets coincide with 258 the top percentile DeepND predicted ASD risk genes (Fisher’s exact test, *P =* 1.14×10^*−*20^) and 99 of its targets coincide with the top percentile DeepND predicted ID risk genes (Fisher’s exact test, *P =* 7.18 × 10^*−*18^). 62 targets are shared among ASD and ID. Note that 204 genes overlap in top percentile ASD and ID risk genes. This means 30% of the shared set of genes among two disorders are targeted by SMARCD1 (Supplementary Table 3). When the union set of the top percentile genes for ASD and ID are considered (312 genes), we find that the top transcription factor targeting these is again SMARCD1 (82 out of 2071 targets; Fisher’s exact test, *P =* 1.24×10^*−*21^). This results indicates that SMARCD1 might be playing role in the convergent etiologies of these comorbid disorders as a shared upstream chromatin modeller.

### 2.4 Other Interactions between ASD and ID Genes

We further investigate the protein-protein interactions (PPI) between the top percentile risk genes in tissue specific PPI network for frontal cortex obtained from DifferentialNet database [6] using the NetworkAnalyst system [85]. This analysis reveals several hub proteins such as HECW2, EP300, and CREBBP1 as shown in Figure 4b. In this list, *HECW*2 gene stands out as it has the highest degree in this network and has very low prior risk for both ASD (TADA *P* = 0.95) and ID (extTADA *Q* = 0.97). Yet, it is in the top percentile for ID risk. Note that DeepND did not use any PPI network information in its reasoning, and yet, was able to identify this hub protein which has been linked with ASD [41] and ID [35] via de novo disruptive mutations in simplex families.

### 2.5 Evaluation of Novel Predictions and Identification of Candidates within ASD and ID Associated CNV Regions

Recurrent copy number variations in several regions of the genome are associated with ASD and ID etiology. However, these are large regions and it is not clear which genes in particular are driver genes. We investigate the DeepND risk ranking of genes within (i) six regions which frequently harbor ASD-related CNVs (16p11.2, 15q11-13, 15q13.3, 1q21.1 and 22q11) [49], and (ii) six regions which were reported to harbor mental retardation related CNVs (16p11.2, 1q21.1, 22q11.21, 22q11.22, 16p13 and 17p11.2) [59]. Note that these CNVs in these regions might confer risk for both disorders (Supplementary Table 4).

DeepND highly ranks several candidate genes for ASD and ID, which (i) are within these CNV regions, (ii) have low prior risk (e.g., E2 or lower, low TADA *P* value etc.), and (iii) low posterior risk assigned by other algorithms (Supplementary Table 4). We discuss some of them below.

*NIPA2* is an ASD E3-E4 gene which encodes a magnesium transporter and located on 15q11-13. It does not participate in any of the relevant gene sets (e.g., risk pathways) and it has a low TADA *P =* 0.768. Although it is ranked in the 5^*th*^ decile for ASD, it is listed in the top decile by DeepND for ID. Its linkage to Prader-Willi Syndrome by [31] also suggest that *NIPA2* might is an important candidate for ID.

DeepND links *MICAL3* to ASD and ID which is in the 22q11.21-22 CNV region. It is a gene related to actin and Rab GTPase binding, ranked in as the 191^*st*^ for ASD risk and 147^*th*^ for ID risk despite having ASD-TADA *Q* = 0.77 and not being in relevant risk gene groups. Krishnan et al. rank it 11, 665^*th*^ and DAMAGES score ranks it 1916^*th*^. DAWN provides no ranking as it is not co-expressed with other is networks of interest for DAWN. While disruption in Rab GTPase cycle was shown to cause intellectual disability [76], there are no established ties between *MICAL3* and ASD/ID. Moreover, MICAL family genes are related to cytoskeletal organization which recently has been deemed as an important molecular function in ASD etiology [75, 65].

It is also an important task to pinpoint genes that might seem related to a neurodevelopmental disorder but actually is not. This also corresponds to untangling the genetic architectures of comorbid disorders. One example that DeepND pinpoints is *ZBTB20*. Although *ZBTB20* is not located within investigated CNV regions, it is regarded as an important risk gene for ASD. It is a CHD8 target and is listed as an E1 gene for ASD with TADA *Q =* 0.17. DAWN and Krishnan et al. rank it as *∼* 500^*th*^ ASD risk gene. However, DeepND consistently ranks it in the last decile with 0.252 probability of being an ASD risk gene. DeepND predicts that there is somewhat a higher chance for *ZBTB20* to be a candidate risk gene for ID while it is still in the 5^*th*^ decile. This gene acts as a transcriptional repressor, it plays a role in various processes such as postnatal growth. Its shown relation to Primrose Syndrome which is specifically characterized by intellectual disability rather than other symptoms of ASD [14, 13] also suggests that *ZBTB20* has a closer relation to ID. Yet, more mutational evidence is required for a more concrete assessment.

Finally, DeepND ranks *LMTK2* as the 2^*nd*^ and 7^*th*^ highest risk gene for ASD and ID, respectively, while other algorithms do not even list it in the top 1000 (Supplementary Table 1). It has very low TADA Q values for both disorders. However, it is a target of CHD8 and FMRP, and also is a post synaptic gene. It is a gene playing role in nerve growth factor (NGF)-TrkA signalling and plays a role in spermatogenesis. With this backgorund in the literature, we think this could be a novel new candidate for both disorders identified by DeepND.

## 3 Discussion

Neurodevelopmental disorders have been challenging geneticist and neuroscientists for decades with complex genetic architectures harboring hundreds of risk genes. Tracing inherited rare and *de novo* variation burden has been the main driver of risk gene discovery. However, overlapping genetic components and confounding clinical phenotypes make it hard to pinpoint disorder-specific susceptible genes and to understand differences. For instance, Satterstrom et al. pinpoint 102 ASD risk genes with FDR *<* 10% with the largest ASD cohort to date covering nearly 6.5k trios [75]. Yet, they still need to manually segregate these risk genes into two groups as (i) 53 ASD predominant risk genes which are distributed across a spectrum of ASD phenotypes, and (ii) 49 neurodevelopmental delay risk genes causing impaired cognitive, social, and motor skills. Thus, comorbidity is a further obstacle to be reckoned with in addition to identifying individual susceptible genes. Nevertheless, the shared risk genes and biological pathways offer opportunities for computational risk assessment methods which were not explored before. So far, only disorder specific analyses were possible by design of the network-based gene discovery algorithms. These are limited in power due to distinct datasets which lead to limited cohort sizes. Here, we proposed a novel approach which can co-analyze comorbid neurodevelopmental disorders for gene risk assessment. The method is able to leverage the shared information using multitask learning paradigm for the first time for this task. DeepND learns both a shared and a disorder-specific set of weights to calculate the genome-wide risk for each disorder. DeepND is a multitask deep learner which uses state-of-the-art techniques underneath such as graph convolution and mixture of experts to learn non-linear relationships of genes on 52 brain co-expression networks. The model is also interpretable as it is able to learn which neurodevelopmental windows (i.e., networks) provide more information for distinguishing high risk genes, and thus, are important for understanding disease etiology. Our benchmarks show these techniques enable DeepND to outperform existing gene discovery methods.

We focus on ASD and ID in this study and identify similarities such as shared affected pathways and neurodevelopmental windows, and differences such as regulatory relationships and novel risk genes. We think the findings in this paper will help guiding neuroscientists researching ASD and ID in prioritizing downstream functional studies. DeepND is not an algorithm specific to these disorders though. It can easily be extended to consider other comorbid disorders such as schizophrenia and epilepsy by adding similar singletask layers for each disorder. It can also be used for other comorbid disorders that are not related to neurodevelopment at all.

We demonstrated the advantage of being able to employ multiple co-expression networks and pointed to cases where, for instance, DAWN was not able to capture relationships between genes as they are limited with a single co-expression network. For the clarity of discussion and interpretability, we focused on only networks produced from BrainSpan dataset. However, DeepND can also employ any combination of other types of gene interaction networks such as protein interaction networks.

## 4 Methods

### 4.1 Ground Truth Risk Gene Sets

Our genome-wide gene-risk prediction algorithm is based on a semi-supervised learning framework in which some of the samples (i.e., genes) are labeled (as ASD/ID risk gene or not) and they are used to learn a ranking for the unlabeled samples.

We obtain the labels for ASD from SFARI dataset [2]. These genes are classified into three evidence levels indicating the quality of the evidence (E1 - E3, E1 indicating the highest risk). The list contains 185 E1 genes (from SFARI Cat I), 200 E2 genes (from SFARI Cat II) and 470 E3 genes (from SFARI Cat III)as positively labeled ASD-risk genes. We also obtain 893 non-mental health related genes from OMIM and Krishnan et al. as negatively labeled genes. The list and corresponding evidence levels are listed in Supplementary Table 1. In the loss calculations during training of the model, genes in these categories are assigned the following weights: E1 genes (1.0), E2 genes (0.50), E3/E4 genes (0.25) and negative genes (1.0). The performance is evaluated only on E1 genes for the sake of compatibility with other methods.

For ID, we rely on review studies from the literature which provide in depth analyses and lists of ID risk genes. We considered the *known gene* lists from 2 landmark review studies as our base ground truth ID risk gene list [26, 34]. We divide this set into 2 parts (i.e., E1 and E2) with respect to evidence obtained from 4 other studies [79, 39, 12, 45]. The genes which are indicated by 5 or more studies are assigned to the highest risk class, E1. Remaining genes which are indicated by 4 studies are assigned to the second highest risk class, E2. See Supplementary Table 2 for a detailed breakdown of evidence for each gene. We use the same set of negative genes as ASD which are non-mental health related genes. Consequently, we have 123 E1, 224 E2 and 232 E3 genes and 893 negative genes for ID. See Supplementary Table 1 for a complete list of ground truth genes for both disorders. The weights we use per gene class are as follows: E1 genes (1.0), E2 genes (0.50), E3 genes (0.25), and negative genes (1.0). For the sake of consistency, we also report the performance of the model on E1 genes for ID.

### 4.2 Gene Risk Features

The only gene risk feature we use the gene pLI for both disorders.See Supplementary Table 1 for pLI of each gene obtained from [75]. Note that neurodevelopmental risk genes are known to be conserved. Yet, the ground truth labels described in Section 4.1 are assigned independently and has no ties to this information.

### 4.3 Gene Co-expression Networks

We used the BrainSpan dataset of the Allen Brain Atlas [78, 44] in order to model gene interactions through neurodevelopment and generated a spatio-temporal system of gene co-expression networks. This dataset contains 57 postmortem brains (16 regions) that span 15 consecutive neurodevelopmental periods from 8 postconception weeks to 40 years. To partition the dataset into developmental periods and clusters of brain regions, we follow the practice in Willsey et al. (2013) [83]. Brain regions were hierarchically clustered according to their transcriptional similarity and four clusters were obtained (Figure 3b): (i) V1C-STC (primary visual cortex and superior temporal cortex), (ii) PFC-MSC (prefrontal cortex and primary motor-somatosensory cortex), (iii) SHA (striatum, hippocampal anlage/hippocampus, and amygladia), and (iv) MDCBC (mediodorsal nucleus of the thalamus and cerebellar cortex). In the temporal dimension, 13 neurodevelopmental windows (Figure 3a) were obtained using a sliding window approach (i.e., [1–3], [3–5], …, [13–15]). A spatio-temporal window of neurodevelopment and its corresponding co-expression network is denoted by the abbreviation for its brain region cluster followed by the time window of interest, e.g. “PFC-MSC(1-3)” represents interactions among genes in the region PFC-MSC during the time interval [1–3].

Using the above-mentioned partitioning, we obtained 52 spatio-temporal gene co-expression networks, each of which contain 25,825 nodes representing genes. An undirected edge between two nodes is created if their absolute Pearson correlation coefficient |*r*| is greater than or equal to 0.8 in the related partition of BrainSpan data.

### 4.4 DeepND Model

#### Problem Formulation

Each 52 co-expression network *j* described in Section 4.3 is represented as a graph *G*_*j*_ = (*V, E*_*j*_), where the vertex set *V =* (*v*_1_, …, *v*_*n*_) contains genes in the human genome and *E* _*j*_ ∈ {0, 1}^*n*×*n*^ denotes the binary adjacency matrix. Note that *n =* 25,825. Let *X*_*D*_ ∈ ℝ ^*n*×*d*^ be the feature matrix for disorder D where each row *X*_*D*_[*i*] is a list of *d* features associated with gene (and node) *i*, ∀*i* ∈ [1, *n*]. Let *y*_*ASD*_ be an *l* dimensional vector, where *y*_*ASD*_[i] = 1 if the the node *i* is a risk gene for ASD, and *y*_*ASD*_[i] = 0 if the gene is non-mental health related, ∀*i* ∈ [1, *l*], *l < n*. Note that, in this semi-supervised learning task, only first *l* genes out of *n* have labels. *y*_*ID*_ is defined similarly for ID, using its ground truth risk and non-risk gene sets. The goal of the algorithm is to learn a function *f* (*X*_*ASD*_, *X*_*ID*_, *y*_*ASD*_, *y*_*ID*_, *G*_1_, …, *G*_52_) → *P* ∈ℝ ^*n*×2^. *P*[*i*][*ASD*] denotes *p*(*y*_*ASD*_[*i*] = 1), and *P*[*i*][*ID*] denotes *p*(*y*_*ID*_[*i*] = 1), ∀*i* ∈ [1, *n*].

#### Graph Convolutional Network Model

Convolutional Neural Networks (CNNs) have revolutionized the computer vision field by significantly improving the state-of-the art by extracting local patterns on grid-structured data [50]. Applying the same principle on arbitrarily structured graph data have also been a success [9, 22, 37]. While all these spectral approaches have proven useful, they are computationally expensive. Kipf and Welling have proposed an approach (graph convolutional network - GCN) to approximate the filters as a Chebyshev expansion of the graph Laplacian [19] and let them operate on the 1-hop neighborhood of each node [48]. This fast and scalable approach extracts a network-adjusted feature vector for each node (i.e., embedding) which incorporates high-dimensional information about each node’s neighborhood in the network. The convolution operation of DeepND is based on this method. Given a gene co-expression network *G*_*j*_, GCN inputs the normalized adjacency matrix *Ê* _*j*_ with self loops (i.e., *Ê*_*j*_ [*i, i*] = 1, ∀*i* ∈ [1, *n*]) and the feature vector *X*_*D*_[*i*] ∈ ℝ^*n×d*^, for gene *i* and for disorder *D. d* = 1 in this application and shared among ASD and ID. Then, the first layer embedding 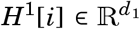 produced by GCN is computed using the following propagation rule 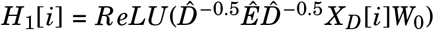 where *W*_0_ is the weight matrix at the input layer to be learnt, *ReLU* is the rectified linear unit function and 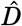 is the normalized version of a diagonal matrix where 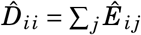. We pick *d*_1_ = 4 in this application. Each subsequent layer *k* is defined similarly as follows: 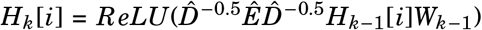. Thus, the output of a *k*-layered GCN, for gene *i* on co-expression network *j* is denoted as follows: 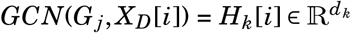. In this application, we use 2 GCN layers and *d*_2_ = 1. The final layer is softmax to produce probabilities for the positive class. That is, the output of a GCN model *j*, for a gene *i* is 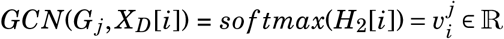.

#### Mixture of Experts Model

Mixture of experts model (MoE) is an ensemble machine learning approach which aims to find out informative models (experts) among a collection [58]. Specifically, MoE inputs the features and assigns weights to the outputs of the experts. In DeepND architecture, individual experts are the 52 GCNs which operate on 52 gene co-expression networks as explained above (*G*_1_, …, *G*_52_). For every gene *i*, MoE inputs *X*_*D*_[*i*] and produces a weight 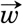 of length 52 which is passed through a softwax layer (i.e., 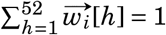). The output of the GCNs are weighted by this network and the weighted sum is used to produce a risk probability for every gene *i* using softmax. That is, the 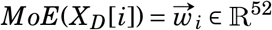. The GCNs produce 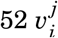 values (one per co-expression network) which are concatenated to produce 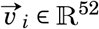. Finally, the following dot product is used to produce the risk probability of gene *i* for disorder 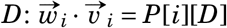.

#### Multitask Learning Model

Above-mentioned GCN and MoE cascade inputs *X*_*D*_[*i*] and co-expression networks *G*_1_, .., *G*_52_ along with labels for a single disorder to predict the risk probability for every gene. Thus, it corresponds to the single-task version of DeepND (i.e., DeepND-ST.) On the other hand, DeepND is designed to work concurrently with multiple disorders (i.e., ASD and ID.) DeepND employs one DeepND-ST cascade per disorder and puts a multi-layer perceptron (MLP) as a precursor to two DeepND-ST subnetworks. The weights learnt on these subnetworks are only affected by the back-propagated loss of the corresponding disorder, and hence, these are single task parts of the architecture. On the contrary, the weights learnt on the MLP part are affected by the loss back-propagated from both subnetworks. Thus, this part corresponds to the multitask component of the DeepND architecture. The MLP layer inputs the union of *X*_*ASD*_ and *X*_*ID*_ and passes it through a fully connected layer followed by Leaky ReLU activation (negative slope = *−*1.5) to learn a weight matrix *W*_*MLP*_ and output a *d*^*′*^ dimensional embedding to be input DeepND-ST instead of *X*_*ASD*_[*i*] and *X*_*ID*_[*i*]. That is, 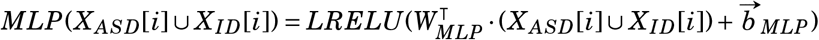.

#### Learning and the Cross Validation Setting of DeepND

To evaluate both DeepND-ST and DeepND approaches, we use a five-fold cross-validation scheme. All labeled genes are uniformly and randomly distributed to the folds. At each training iteration, we leave one fold for validation and one fold for testing. We train the model on the remaining three folds of the labeled genes. For all the genes in the left-out fold, their feature vectors are nullified when input to training in order to prevent information leakage.

The model is trained up to 1000 epochs with early stop which is determined with respect to the loss calculated on the validation fold using only E1 genes as the positives and all negative genes. Once the model converges, the test performance is reported on the test fold in the same manner as in the validation fold. The model uses cross entropy loss (evidence weighted) and ADAM optimizer [47] for updating the weights which are initialized using Xavier initialization [29]. DeepND-ST uses a fixed learning rate of 7 × 10^*−*4^ for both disorders. DeepND uses the learning rates of 7 × 10^−4^, 7 × 10^−4^ and 7 × 10^−3^, for the shared layer, ASD single task layer, and ID single task layer, respectively. To ensure proper convergence of DeepND, should one of the singletask subnetworks converge, the learning rate of that subnetwork and the shared layer are cut down twenty folds. This lets the yet underfit subnetwork to keep learning till early stop or the epoch limit.

Above-mentioned procedure produces 4 results as for every left-out test fold as all remaining 4 folds are used as validation. We repeat this process 10 times with random initialization to obtain a total of 200 results for performance comparison (Figure 2). For ASD, positively labeled genes’ weights equal to 1.00 (E1), 0.50 (E2), 0.25 (E3) and 0.25 (E4). For ID, positively labeled genes’ weights equal to 1.00 (E1) and 0.50 (E2). For both disorders, negatively labeled genes have weight 1.00. Note that this procedure is in line with the setting of Krishnan et al. for fairness of comparison.

### 4.5 Genome-wide risk prediction for ASD and ID and Comparison with Other Methods

We compare the performance of DeepND-ST and DeepND with other state-of-the-art network based neurodevelopmental disorder gene risk assessment methods from the literature which output a genome-wide risk ranking: Krishnan et al. [49], DAWN [55], DAMAGES score [84], Random Forest based method of Duda et al. [21] and forecASD [8].

We run Krishnan et al.’s approach as described in their manuscript [49]. This is an evidence-weighted Support Vector Machine (SVM) classifier which identifies risk genes based on similarity of network features. They use a human brain-specific functional interaction network to generate features as input [32]. Note that, the ground truth gene set is the same as ours as well as the evidence weights for ASD. We perform a 5-fold cross validation. That is, for each iteration, we train their SVM model on 80% of the labeled genes and evaluate the model on E1 genes and all negative genes in the left-out 20% of the labeled genes as suggested. We repeat this procedure 10 times. We post-process SVM outputs to produce risk probabilities using isotonic regression which ensures that the gene ranking is preserved. We use the pLI value of each gene as the dependent variable. In a 10-fold cross validation setting, we detect *knots* on the left-out fold, and fit another isotonic regression line to interpolate the knots. We use SVM output for all genes to produce a gene risk probability values for the corresponding disorder and produce the genome-wide risk ranking. We compare this method and DeepND-ST/DeepND with respect to (i) the area under receiver operating curve (AUC) and area under precision recall curve (AUPR) distributions calculated on the left-out fold at each cross validation iteration (Figure 2a-b), (ii) Matthews Correlation Coefficient of the final predictions with the ground truth (using a rank percentage threshold - Figure 2c), and (iii) precision-recall curves (Figure 2d).

DAWN is a hidden Markov random field based approach that assigns a posterior, network-adjusted disorder risk score to every gene based on guilt by association principle. It inputs TADA p-values as prior features along with a partial co-expression network to assess connectivity. We input the TADA p-values to DAWN [75, 63]. The method also uses partial co-expression networks. We use two networks which the authors suggest as the most useful for this task in their manuscript [55]. These represent prefrontal cortex/mid-fetal period (i.e., PFC-MSC 3-5 and PFC-MSC 4-6). We generate these networks using the RNA-Seq data in the BrainSpan dataset. Note that DeepND utilizes the same dataset and uses these networks and 50 others. Instead of partial co-expression networks, DeepND uses co-expression networks. The DAMAGES score is a principal component analysis based technique that assess the risk of genes based on (i) the similarity of their expression profiles in 24 specific mouse central nervous system cell types, (ii) the enrichment of mutations in cases as opposed to controls, and (iii) the pLI score of the gene. We directly obtain the risk ranking from Shen and Zhang, 2017 [84]. We also benchmark the Random Forest based method presented by Duda et al. as described in [21]. The method inputs a combination of (i) gene expression data from several microarray studies, (ii) PPI interactions from multiple databases, (iii) a quantitative physical interaction (PI) score for all protein isoform pairs in mouse, and (iv) phenotype annotations from the Mouse Genome Informatics (MGI) database. They combine these modalities by a Bayesian network and the final probability of functional interaction between gene pairs are represented with a matrix. Since this method does not support evidence weights for the ground truth set, in the experimental setting, we used our ground truth gene set for ASD after discarding the evidence information to obtain results. We compare all above mentioned methods with respect to (i) Matthews Correlation Coefficient of the final predictions with the ground truth (using a rank percentage threshold - Figure 2c), and (ii) precision-recall curves (Figure 2d).

Apart from the previously mentioned methods, we also consider forecASD which is a random forest based ensemble method. It inputs features derived from (i) BrainSpan transcriptome, (ii) the STRING PPI network and the outputs of several previously published ASD gene prediction methods: Krishnan et al. [49], DAWN [55], DAMAGES score [84] and TADA [74]. These features are combined in a two-layer architecture. In the first layer, each derived feature set is used to train a random forest branch. Then, in the second step, the results from the first layer are combined and are used to train the final ensemble classifier. We trained the original forecASD architecture with our ground truth sets for ASD and ID and obtained their results. Then, in a separate run, we also added the final ranking of DeepND as an extra feature and retrained the model. We compared these two results with our final ranking using (i) Matthews Correlation Coefficient of the final predictions with the ground truth (using a rank percentage threshold), and (ii) precision-recall curves (Supplementary Figure 1). All algorithms are trained and tested on a SuperMicro SuperServer 4029GP-TRT with 2 Intel Xeon Gold 6140 Processors (2.3GHz, 24.75M cache), 251GB RAM, 6 NVIDIA GeForce RTX 2080 Ti (11GB, 352Bit) and 2 NVIDIA TITAN RTX GPUs (24GB, 384Bit). For DeepND-ST, we used 3 2080 RTX and 1 TITAN RTX cards and the cross validation setup took approximately 5 hours. For DeepND, we used 5 RTX 2080 and 1 TITAN RTX cards and the cross validation setup took approximately 16 hours.

### 4.6 Enrichment Analyses

We evaluate the enrichment of DeepND’s ASD and ID gene risk rankings in gene lists which are known to be enriched in disorder-risk genes in the literature. These lists are (i) targets of transcription regulators like CHD8 [15], FMRP [16, 72], RBFOX1 [81, 82] and TOP1 [46]; (ii) Susceptibility pathways like WNT signaling [43] and MAPK signaling [68]; and (iii) affected biological processes and molecular functions like, post-synaptic density complex [7, 87], histone modification [18, 42]). We use the binomial test to determine whether the top decile in the corresponding ranking significantly deviate from uniform enrichment (Figure 5).

We perform a Gene Ontology term enrichment analysis of the top percentile predictions using the Enrichr system [11, 51] with respect to Biological Process terms and Molecular Function terms. As both analyses point to *transcription factor regulation*, we investigate the connectivity of the high risk genes in the ChEA Database [52] which is a large repository that lists experimentally validated transcriptional regulation relationships in various organisms.

### 4.7 Spatio-Temporal Network Analyses

We investigate which GCNs (i.e., neurodevelopmental windows) are better predictors of gene risk for each disorder. For each gene *i*, the average risk probability assigned by the corresponding GCN is calculated over all iterations such that *i* is in the test fold. The average values for top percentile genes for each disorder are shown in Figure 3.

To check whether obtained average risk probabilities per neurodevelopmental window that are assigned by that GCN are empirically significant, we performed a permutation test. First, we randomly redistribute the labels of the ground truth genes, while keeping the counts same. We obtain 100 randomized ground truth sets. Keeping all other settings same, we train 100 DeepND models and obtain 100 ASD and 100 ID risk rankings. Using these rankings, we calculate the the average risk probability assigned by each GCN and obtain two background distributions for each GCN; one for ASD and one for ID (Supplementary Figures 4 and 5). We then compare the original average probability assigned by that GCN with the corresponding background distribution to assess significance.

## Code Availability

DeepND is implemented and released at http://github.com/ciceklab/deepnd. We provide the environment which contains all dependencies for an easy setup. We give a small example to train and test both DeepND-ST/ DeepND models. Finally, we provide the code and links to the full set of datasets to reproduce the results (genomewide risk rankings and data for heatmaps) presented in this manuscript for ASD and ID.

## Data Availability

All datasets used in this study are publicly available, which are referenced in the relevant methods subsections. The Genotype-Tissue Expression (GTEx) Project was supported by the Common Fund of the Office of the Director of the National Institutes of Health, and by NCI, NHGRI, NHLBI, NIDA, NIMH, and NINDS. The GTEx data used for the analyses described in this manuscript were obtained from the GTEx Portal on Feb 2020. All data supporting the key findings such as gene risk evidence levels and gene risk predictions are available within the article and corresponding supplementary tables.

## Acknowledgements

We thank all families who participated in the studies which enable this study with data. Authors would like to acknowledge Autism Sequencing Consortium PIs and contributors. This work was supported by a pilot grant from the Simons Foundation (SFARI 640935, AEC). We also acknowledge the support by TUBA GEBIP 2017 and Bilim Akademisi BAGEP 2020 Awards to AEC.

## Author Contributions

AEC conceived, designed, and supervised the study. IB and OK implemented the DeepND software. IB, OK and AEC performed the computational experiments and wrote the manuscript.

## Supplementary Figure and Table Legends

***Supplementary Figure 1***. Figure compares forecASD, DeepND and forecASD + DeepND performances. (a) MCC between the rankings of each method and the ground truth genes are shown for varying rank percentage threshold values (x axis). This ranking threshold *p* sets the top *p*% of a ranking as positive predictions. and (b) precision-recall curves.

***Supplementary Figure 2***. (a) The percentage overlap between corresponding deciles of the ASD and ID genome-wide risk rankings are shown. (b) The percentage overlap between corresponding percentiles of the ASD and ID genome-wide risk rankings are shown.

***Supplementary Figure 3***. Heatmaps show which spatio-temporal windows lead to assignment of higher risk probabilities to the top percentile genes for ASD (left) and ID (right). The numbers in boxes for network probabilities are softmaxed outputs of each respective GCN, averaged for top percentile genes and then normalized. The categorization of brain regions and time windows are provided in Figure 3.

***Supplementary Figure 4***. Results for the permutation test to assess the significance of the informativeness of each neurodevelopmental window for ASD. The boxplots show the distribution of the average probabilities assigned by the corresponding GCN to top percentile genes, for each of the 100 randomized ground truth sets. The red cross indicates the result obtained with actual ground truth labels and match with the results presented in Figure 3b.

***Supplementary Figure 5***. Results for the permutation test to assess the significance of the informativeness of each neurodevelopmental window for ID. The boxplots show the distribution of the average probabilities assigned by the corresponding GCN to top percentile genes, for each of the 100 randomized ground truth sets. The red cross indicates the result obtained with actual ground truth labels and match with the results presented in Figure 3b.

***Supplementary Table 1***. Genome-wide risk probability predictions and rankings of DeepND for ASD and ID. The table marks the gold standard genes: 855 positively and 893 negatively labeled genes for ASD; and 579 positively and 893 negatively labeled genes for ID are given along with their evidence levels (E1 - E4, E1 indicating the highest risk). The table also provides the rankings from other gene discovery algorithms from the literature. For each gene, participation in disorder-related gene sets are provided (e.g., WNT pathway, *CHD8* targets etc.). We also provide 29 gene specific details compiled from the literature.

***Supplementary Table 2***. The table provides information about the studies used to generate the ground truth labels for ID. For each gene, the base studies which indicates it as a risk gene are provided.

***Supplementary Table 3***. The table lists (i) the GO enrichment analysis for the top percentile risk genes for both disorders (Biological Process and Molecular Function). It also provides lists of top transcription factor (TF) regulators whose targets are enriched with the top percentile ASD and/or ID risk genes, respectively. The TF enrichment results are based on the experimentally validated TF-gene relationships in ChEA 2016 database. All results are obtained using the EnrichR system.

***Supplementary Table 4***. DeepND risk rankings for the genes within 6 ASD related (16p11.2, 15q11-13, 15q13.3, 1q21.1 and 22q11) (Krishnan et al., 2016) and 6 mental retardation related CNV regions (16p11.2, 1q21.1, 22q11.21, 22q11.22, 16p13 and 17p11.2). Frequency of these CNV regions within ASD and ID individuals are provided when available.

