## Supplementary figures and images for "Deep multitask learning of gene risk for comorbid neurodevelopmental disorders"

### Supplementary Figure 1

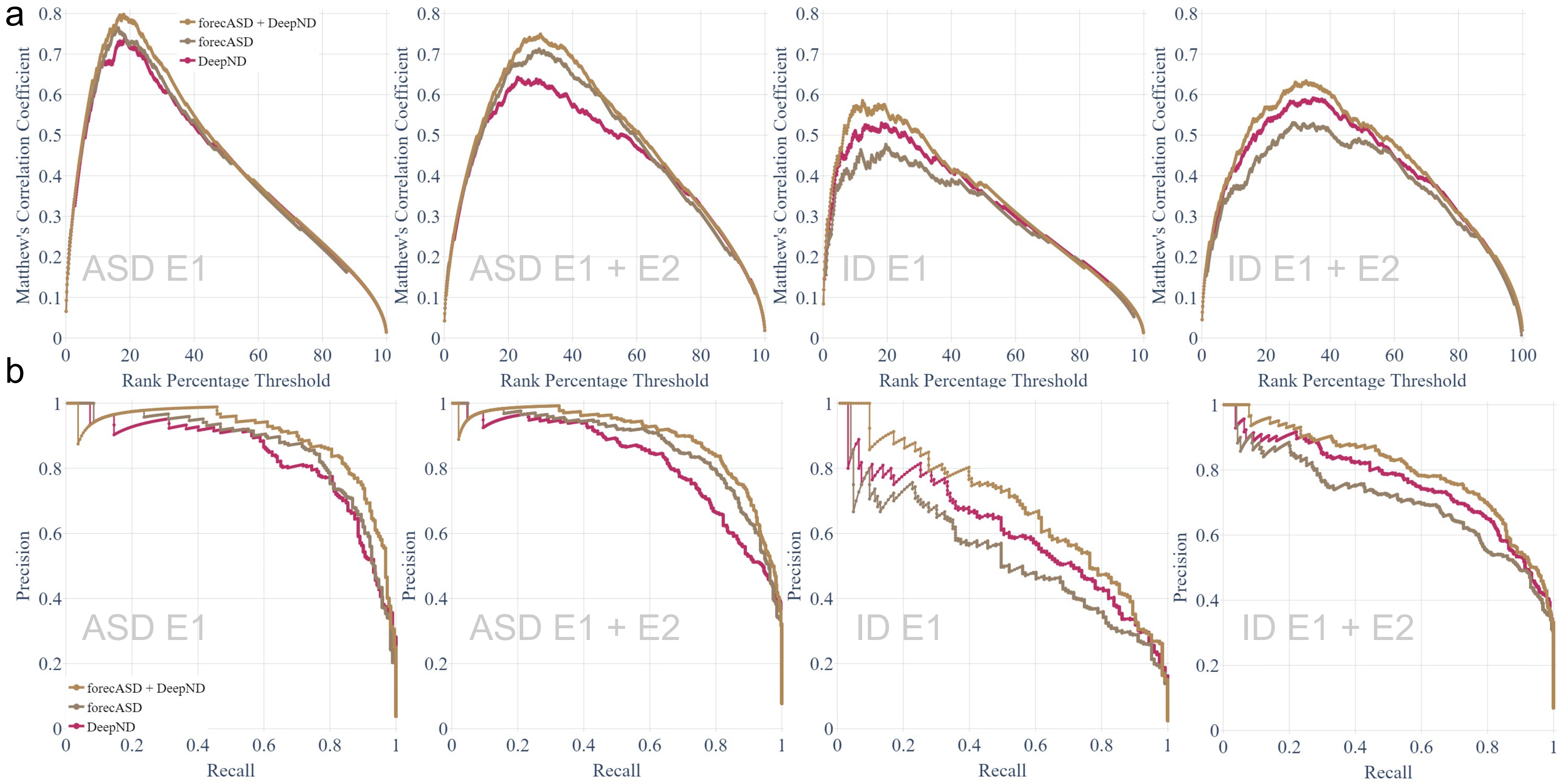

### Supplementary Figure 2

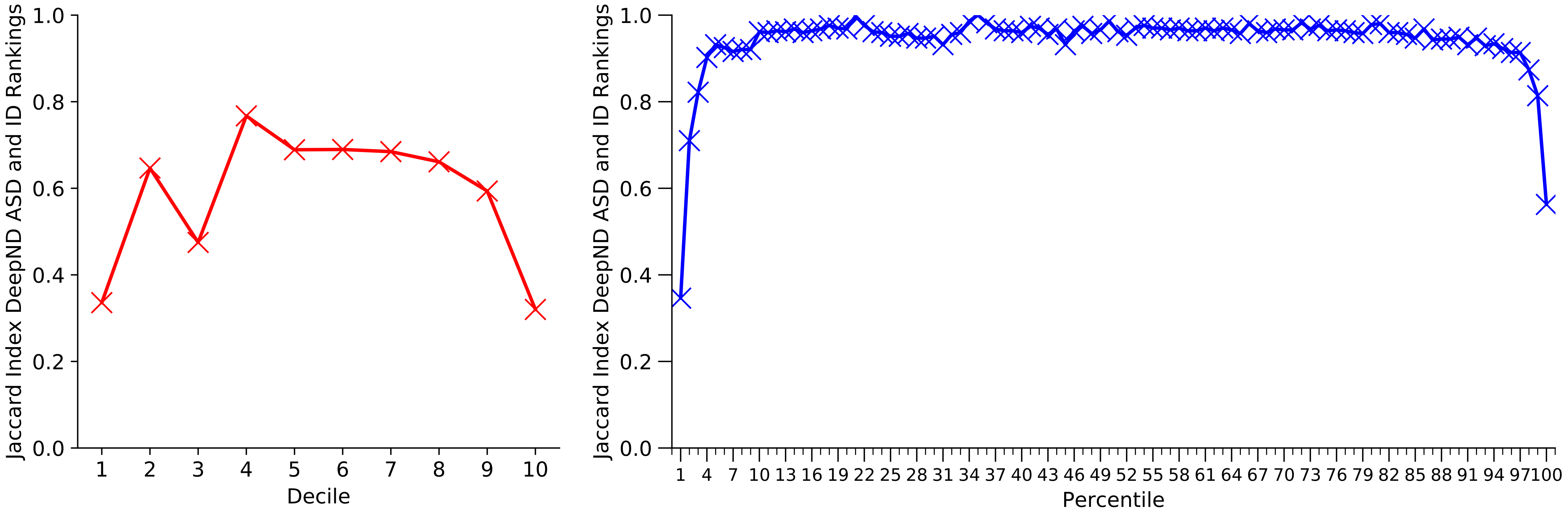

### Supplementary Figure 3

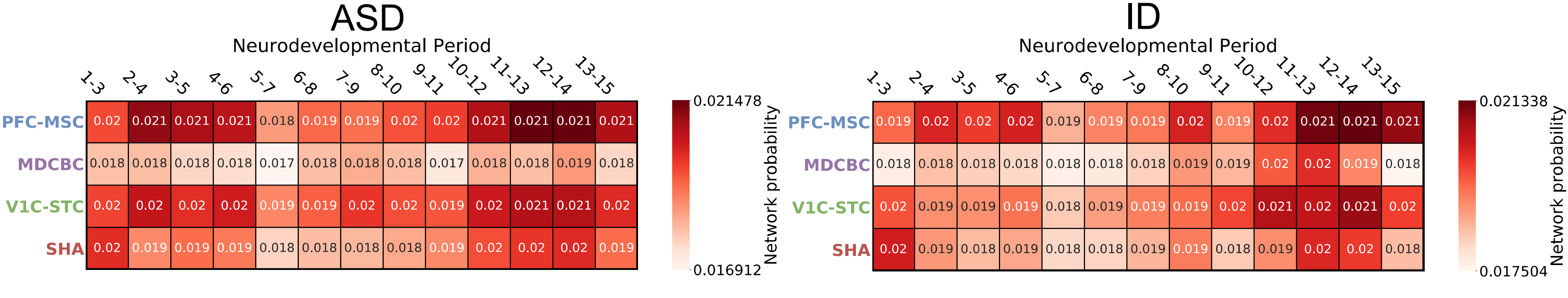

### Supplementary Figure 4

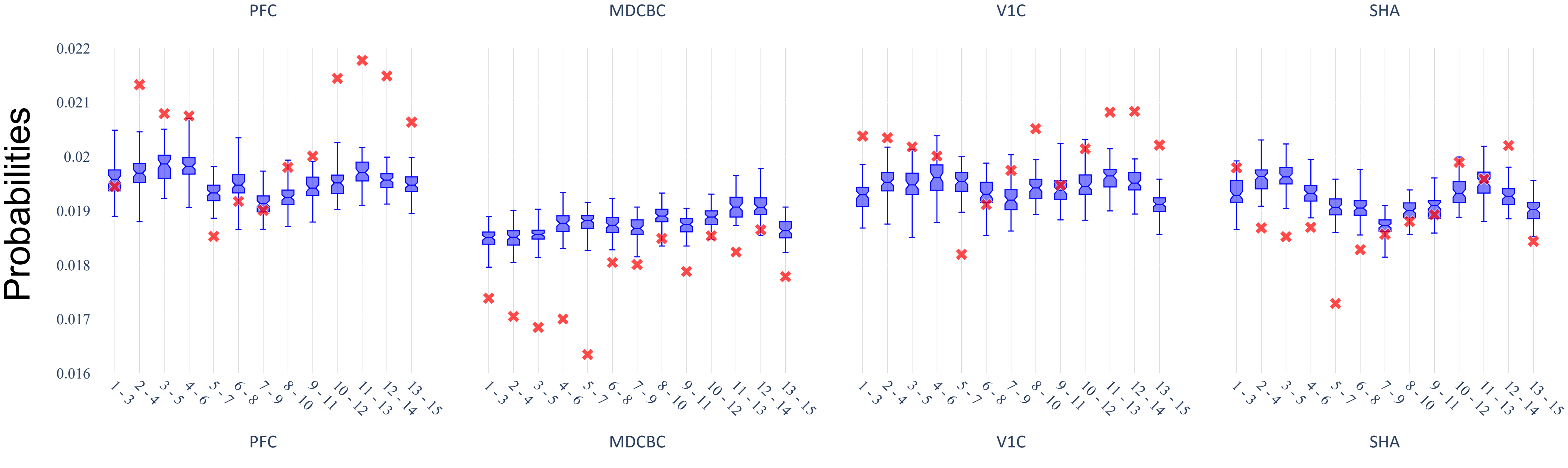

### Supplementary Figure 5

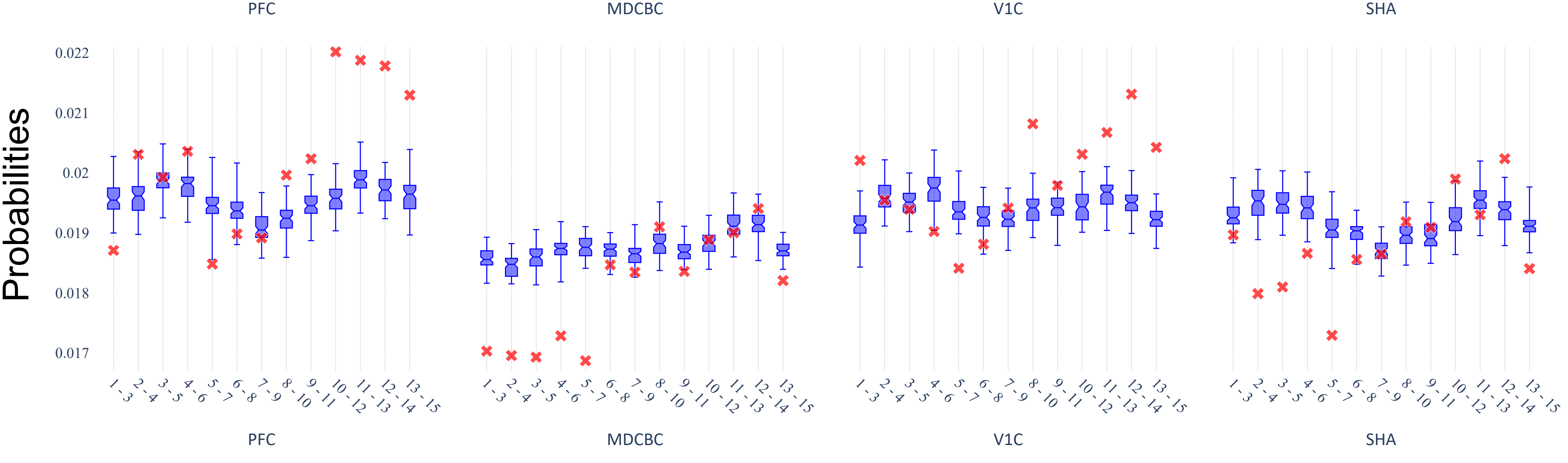
